# Variant- and Vaccination-Specific Alternative Splicing Profiles in SARS-CoV-2 Infections

**DOI:** 10.1101/2023.11.24.568603

**Authors:** Sung-Gwon Lee, Priscilla A. Furth, Lothar Hennighausen, Hye Kyung Lee

## Abstract

The COVID-19 pandemic, caused by the coronavirus SARS-CoV-2, and its subsequent variants has underscored the importance of understanding the host-viral molecular interactions to devise effective therapeutic strategies. A significant aspect of these interactions is the role of alternative splicing in modulating host responses and viral replication mechanisms. Our study sought to delineate the patterns of alternative splicing of RNAs from immune cells across different SARS-CoV-2 variants and vaccination statuses, utilizing a robust dataset of 190 RNA-seq samples from our previous studies, encompassing an average of 212 million reads per sample. We identified a dynamic alteration in alternative splicing and genes related to RNA splicing were highly deactivated in COVID-19 patients and showed variant- and vaccination-specific expression profiles. Overall, Omicron-infected patients exhibited a gene expression profile akin to healthy controls, unlike the Alpha or Beta variants. However, significantly, we found identified a subset of infected individuals, most pronounced in vaccinated patients infected with Omicron variant, that exhibited a specific dynamic in their alternative splicing patterns that was not widely shared amongst the other groups. Our findings underscore the complex interplay between SARS-CoV-2 variants, vaccination-induced immune responses, and alternative splicing, emphasizing the necessity for further investigations into these molecular cross-talks to foster deeper understanding and guide strategic therapeutic development.

## Introduction

The emergence of the coronavirus SARS-CoV-2, the causative agent of COVID-19, has precipitated a global public health crisis^1–3^. The SARS-CoV-2 infection manifests a broad spectrum of symptoms ranging from mild respiratory discomfort to severe acute respiratory syndrome, with potential long-term sequela^4,5^. The quest to unravel the molecular mechanisms of SARS-CoV-2 infection and propagation has burgeoned into genomic, transcriptomic, and proteomic dimensions^6–8^.

Alternative splicing is a cellular process that increases the number of mRNA isoforms and can augments the proteomic diversity and function as well as compromising protein expression through loss of open reading frames and switching from coding to non-coding transcripts ^9–11^. It has been implicated in various diseases including viral infections, where it is poised to play a crucial role in modulating host immune responses and viral replication mechanisms^12–15^. This is also true of SARS-CoV-2 infection^11,15–18^. Specific alternative spliced transcripts present with SARS-CoV-2 infection can lead to reduced antiviral immunity. Examples of alternatively spliced genes that have been previously identified include *CD74* and *LRRFIP1*^15^, and *OAS1*^18^. A specific SARS-CoV-2 protein, NSP16, has been shown to bind to the U1 and U2 splicing RNAs^16^. In short, alternative splicing appears to be pivotal molecular biology realm contributing to the pathophysiology of SARS-CoV-2 infection. But, despite the research directed towards understanding feature of SARS-CoV-2 variants, alteration of immune response induced by infection and vaccination, and genome-wide transcriptome alterations^19–24^, there remains a conspicuous research gap persists concerning the alternative splicing profiles and the splicing machinery that can be affected by variants and vaccination statuses.

Here, we analyzed 190 RNA-seq data sets from five COVID-19 cohorts across four variants infected patients (Alpha, Beta, Gamma, and Omicron) and healthy controls to identify their transcriptome profiles including alternative splicing and gene expression. We also investigated variant- and vaccination-specific transcriptional regulations. This examination allowed us to not only explore the intricate transcriptional landscape underpinning the infection dynamics of different SARS-CoV-2 variants but also the potential modulatory impact of vaccination on host transcriptome.

## Results

### Landscape of splicing across COVID-19 patients infected with four SARS-CoV-2 variants reveals an aberrant global alternative splicing pattern

To investigate the impacts of SARS-CoV-2 variants on alternative splicing of cellular RNAs, we analyzed 190 RNA-seq data from buffy coats of COVID-19 patients infected with four different variants (Alpha, Beta, Gamma, and Omicron), as well as healthy controls (HC) (Table S1). We specifically focused on data from patients within one week of infection to investigate transcriptional changes at an early stage. In total, approximately 40.3 billion reads were mapped to the human genome, achieving an average alignment rate of 95.6%. Using the rMATs^25^, we identified 444,167 alternative splicing events and estimated their exon inclusion levels. Principal component analysis (PCA) showed intermingled profiles of most samples except for Omicron-infected patients (Fig. 1A). A total of 3,381 differential alternative splicing events (DASEs) spanning five distinct alternative splicing categories were identified in COVID-19 patients compared to HC (Fig. 1B). We observed 2,245 DASEs in Alpha-infected patients, while dozens of DASEs were found in Beta- and Gamma-infected patients. Interestingly, 11,996 DASEs were specifically identified in Omicron-infected patients. This finding suggested that the alternative splicing is globally modified in COVID-19 patients.

**Figure 1.**
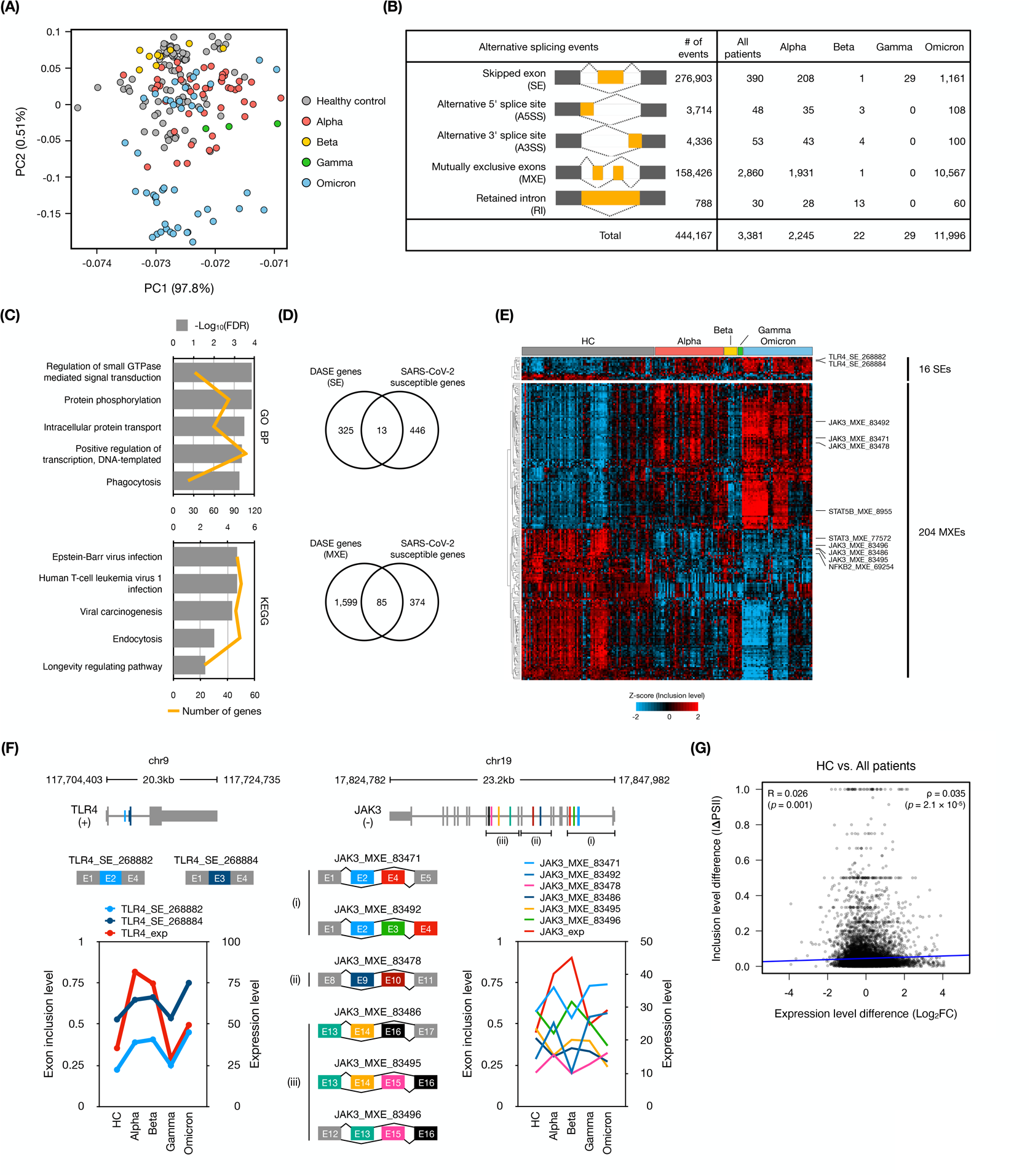
SARS-CoV-2 infection leads to aberrant global alternative splicing. **(A)** Principal component analysis of exon inclusion level from 190 samples. **(B)** Number of differential alternative splicing events (DASEs) of five alternative splicing types. The DASEs were defined as following criteria: absolute PSI differences value > 0.1 and corrected *p*-value < 0.05. **(C)** Results of the top 5 terms of GO biological process and KEGG pathways enriched with the 1,928 genes from 3,381 DASEs (HC vs. All patients). **(D)** Venn diagrams displaying the gene overlap between DASEs in the SE and MXE categories, respectively, and the set of SARS-CoV-2 susceptible genes. **(E)** Heatmap of differentially alternative spliced SARS-CoV-2 susceptible genes between HC and COVID-19 patients. Z-score indicate relative exon inclusion levels. Hierarchical clustering of DASEs was performed with Euclidean distance matrix of relative exon inclusion levels. **(F)** Significant differential spliced events of *TLR4* (chr9:117,704,403-117,724,735) and *JAK3* (chr19:17,824,782-17,847,982) showing two skipped exon (SE) events and six mutually exclusive exon (MXE) events, respectively. Exon inclusion levels represent the usage of spliced exons in the case of SE events, while in the context of MXE events, they indicate the ratio of the second mutually exclusive exon within each event. The expression level refers to the overall gene expression level. The error bar indicates standard deviation of the mean. **(G)** Correlation analysis of expression differences between the inclusion level differences of total genes revealed no significant correlation.

The 1,928 genes from 3,381 DASEs were significantly enriched in GO terms of general cellular functions including regulation of small GTPase mediated signal transduction (GO:0051056; FDR = 1.4 ✕ 10^-4^), intra cellular protein transport (GO:0006886; FDR = 3.3 ✕ 10^-4^) and transcription (GO:0045893; FDR = 4.1 ✕ 10^-4^) (Fig. 1C). Moreover, they are mainly involved in virus infection pathways including Epstein-Barr virus (hsa05169; FDR = 2.0 ✕ 10^-5^) and human T-cell leukemia virus 1 (hsa05166; FDR = 2.0 ✕ 10^-5^). Indeed, these findings suggest that the genes exhibiting altered alternative splicing patterns during SARS-CoV-2 infection play a pivotal role in diverse cellular functions. Moreover, their pronounced involvement in virus infection pathways underscores their significance in shaping the observed alternative splicing patterns.

We compared the predominant alternative splicing events in our results, which comprise approximately 96% of the DASEs, specifically skipped exon (SE) and mutually exclusive exons (MXE) DASEs, with genes associated with susceptibility to viral infections, including SARS-CoV-2 and other viral infections. 13 genes (SE) and 85 genes (MXE) that exhibited associations with viral susceptibility were identified (Fig. 1D). Those genes included 16 SEs and 204 MXEs, showing dynamic splicing alterations observed in COVID-19 patients (Fig. 1E). Among DASEs, we found alternative exon usages of genes related to Janus kinase (JAK) signaling pathway and Toll-like receptor 4 (TLR4) (Fig. 1F). Specifically, exon 2 and 3 of TLR4 displayed higher inclusion levels in COVID-19 patients. Notably, these inclusion patterns have been previously reported in splice variants induced by lipopolysaccharide (LPS) treatment^26,27^. The JAK signaling pathway plays a crucial role in virus infections, including those involving SARS-CoV-2^28^. In our results, JAK3 exhibited six novel MXE events spanning ten exons, indicating a wide range of alternative variants being generated. This underscores the complexity of alternative splicing of TLR4 and JAK3 and its potential significance in the context of viral infections.

In prior research, it was reported that transcription rate significantly influences splicing fidelity in yeast^29^. To investigate the relationship between expression level of genes and their splicing rates in the context of SARS-CoV-2 infection, we compared the transcript level difference and the absolute mean inclusion level difference between HC and COVID-19 patients. However, contrary to our expectations, interrelation between alternative splicing rates and gene expression levels was not observed (Fig. 1G). This outcome suggests that the interplay between gene expression alterations and alternative splicing rates in SARS-CoV-2 infection may differ from what has been observed in yeast, emphasizing the complexity and biological diversity of the alternative splicing process in this context.

### Dysregulated alternative splicing related genes in COVID-19 patients

In our quest to identify genes that might influence alternative splicing changes in COVID-19, we examined global gene expression differences between HC and COVID-19 patients. PCA results revealed pronounced gene expression differences primarily in Alpha and Beta-infected patients, while relatively minor differences observed in Omicron-infected patients compared to HC (Fig. 2A, Fig. S1). We identified 7,529 genes that were differentially expressed (DEGs) in COVID-19 patients. Subsequent ShinyGO analysis associated these DEGs with Coronavirus and Herpes simplex virus infections (Fig. 2B). Interestingly, DEGs were notably enriched in spliceosome-related genes, suggesting dysregulation of spliceosome-related gene expression in COVID-19 patients. Independent DAVID analysis also indicated significant enrichment of 92 DEGs in RNA splicing (GO:0008380), with the majority, 81.5% (75 out of 92 DEGs), being significantly down-regulated in COVID-19 patients (Fig. 2C). Furthermore, we observed significantly reduced expression of hub genes such as HNRNPA1, SNRPA1, SNRPD2, SFPQ, SNRPF, and TARDBP, which interact intensively with other proteins in the protein-protein interaction network (Fig. 2D, E). These findings collectively suggest that alternative splicing machinery is compromised in COVID-19 patients and is closely associated with abnormal global alternative splicing patterns.

**Figure 2.**
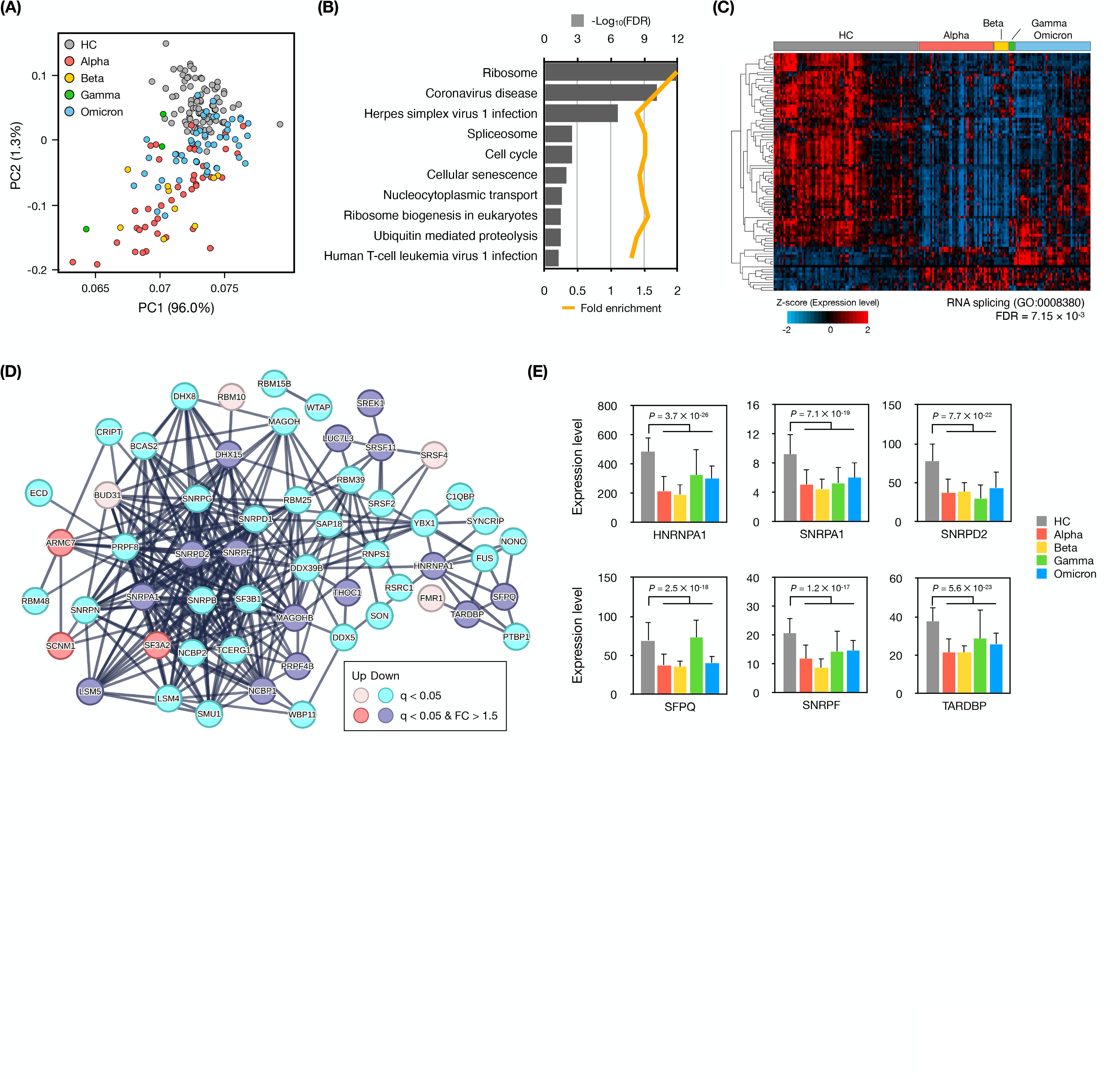
Dysregulated alternative splicing related genes in COVID-19 patients. **(A)** Principal component analysis of gene expression levels from 190 samples. **(B)** Results of the top 10 pathways enriched with the 7,529 DEGs. **(C)** Heatmap of 92 DEGs involved in RNA splicing (GO:0008380). Z-score indicate relative gene expression levels. Hierarchical clustering of DEGs was performed with Euclidean distance matrix of relative gene expression levels. **(D)** Protein-protein network of significant DEGs related to RNA splicing. The color of each gene indicates statistically significant and fold change to HC. **(E)** Gene expression levels of six hub genes which are depressed in COVID-19 patients. The error bar indicates standard deviation of the mean.

### Different regulation of alternative splicing in T and B cell receptor signaling pathways in vaccinated and unvaccinated group of Omicron infected patients

We also observed distinct alternative splicing profiles among patients infected with the Omicron variant (Fig. 1A). Through unsupervised clustering, we delineated two distinct groups within the entire patient cohort. Group 1 consisted of 163 samples (comprising HC, Alpha, Beta, and some Omicron cases), while group 2 comprised the remaining 27 Omicron-infected patients. Notably, we found that group 1 was enriched with unvaccinated individuals, while group 2 predominantly consisted of vaccinated patients (Fig. 3A). The difference in vaccination status between these groups was statistically significant (Fig. 3B).

**Figure 3.**
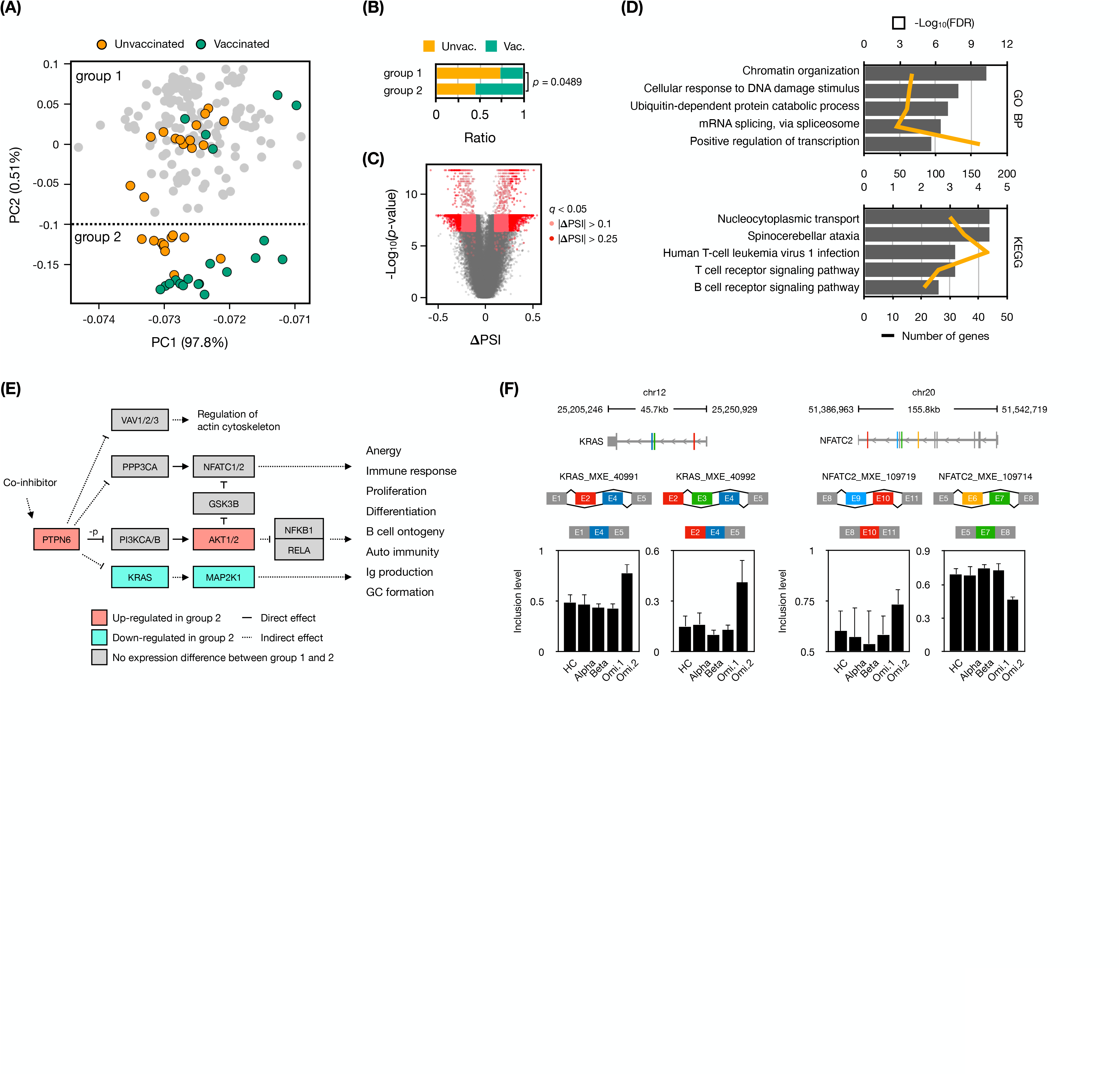
Among Omicron-infected patients, different regulation of alternative splicing in T and B cell receptor signaling related genes was observed between vaccinated and unvaccinated groups. **(A)** PCA of exon inclusion levels from 190 samples, as presented in Figure 1A. The samples are marked differentially based on the vaccination status of Omicron-infected patients. Group 1 and 2 represent clusters established through k-means clustering, as described in Figure SX. **(B)** Individual Omicron-infected patients’ vaccination status ratio within group 1 and group 2. A chi-squared test was conducted to confirm a significant difference in vaccination status proportions between the two groups. **(C)** Volcano plot showing the DASEs between Omicron groups. The x- and y-axis indicate inclusion level difference (𝚫PSI) and negative log_10_ transformed *p*-value. The *q*-value indicated corrected *p*-value. **(D)** Results of the top 5 terms of GO biological process and KEGG pathways enriched with the 1,789 genes from 2964 DASEs (Omi.group1 vs. Omi.group2). **(E)** A subset of shared pathways based on 16 common DASEs identified in the T cell receptor signaling pathway (hsa04660) and the B cell receptor signaling pathway (hsa04662). The coloration of each gene box signifies the expression difference between the two groups, with solid lines indicating a direct effect and dashed lines representing an indirect effect. **(F)** Significant differential spliced events of KRAS (chr12:25,205,246-25,250,929) and NFATC2 (chr20:51,386,963-51,542,719) showing two MXE events, respectively. Inclusion levels represent the usage of spliced exons. The error bar indicates standard deviation of the mean.

We examined the alternative splicing events that changed between these two Omicron-infected patient groups and a total of 19,732 DASEs with the majority being of the MXE type were identified. With a more stringent criterion (an inclusion level difference > 0.25), we identified 2,964 significant DASEs between Omicron group 1 and Omicron group 2 (Fig. 3C). These DASEs were associated with 1,789 genes primarily related to general functions such as chromatin organization (GO:0006325; FDR = 5.4 ✕ 10^-11^), DNA damage response (GO:0006974; FDR = 1.2 ✕ 10^-8^), splicing (GO:0000398; FDR = 3.5 ✕ 10^-7^), transcription (GO:0045944; FDR = 2.1 ✕ 10^-6^), and were closely linked not only to virus infection (hsa05166; FDR = 6.2 ✕ 10^-4^) but also to the T (hsa04660; FDR = 6.2 ✕ 10^-4^) and B cell receptor signaling pathways (hsa04662; FDR = 2.5 ✕ 10^-3^) (Fig. 3D).

Furthermore, we identified 16 common genes that are downstream of the T and B cell receptor signaling pathways. These genes, acting as intermediary genes, regulate immune processes including immune response, proliferation, and differentiation. Interestingly, most of these genes were regulated through splicing without exhibiting expression differences between the groups. Among them, KRAS was identified to have a higher inclusion of exon 4 in Omicron group 2. The isoform with exon 4 inclusion is reported as *KRAS4A* directly regulate glycolysis and apoptosis promotion and is predominantly expressed in endodermal organs^30–32^. Another gene, NFATC2, showed an increased inclusion of exon 10 in Omicron group 2. The isoform with this inclusion has been reported to be ubiquitously expressed across various tissues^33^, however, the decrease in exon 7 inclusion observed in Omicron group 2 represents a novel isoform which has not been reported previously. This implies that, depending on vaccination status, Omicron infection leads to differential regulation of alternative splicing in genes associated with both general cellular functions and immune cell signaling genes, reflecting broader changes in cellular function and immune response.

### Differential regulation of RNA splicing machinery genes across SARS-CoV-2 variants

To identify genes within the RNA splicing machinery that impact the abnormal regulation of alternative splicing in COVID-19 patients, we collected 304 genes related to RNA splicing genes with an FPKM exceeding 5 in at least one sample and conducted a PCA based on their expression values (Fig. 4A). As a result, clear distinctions were made between HC and each patient group, while Alpha and Beta displayed similar trends. Moreover, a tendency of division based on vaccination status was observed within Omicron patients. This suggests that RNA splicing genes are regulated differently among patients infected with respective variants. Upon identifying the top 25 genes contributing significantly to the principal components distinguishing each group, it was observed that 11 genes contributing to HC were expressed at lower levels across all patient groups (Fig. 4B). Additionally, five genes highly contributing to the Alpha and Beta groups exhibited group-specific elevated expression. Furthermore, nine genes in Omicron patients showed distinctively higher expression, with a notably high expression observed particularly in Omicron group 2. Through this, we deduce that the regulation of RNA splicing machinery genes may be associated with the variant-specific host responses to SARS-CoV-2 infection. The differential expression and regulatory patterns of these splicing machinery genes might play pivotal roles in the abnormal alternative splicing regulation observed in COVID-19 patients.

**Figure 4.**
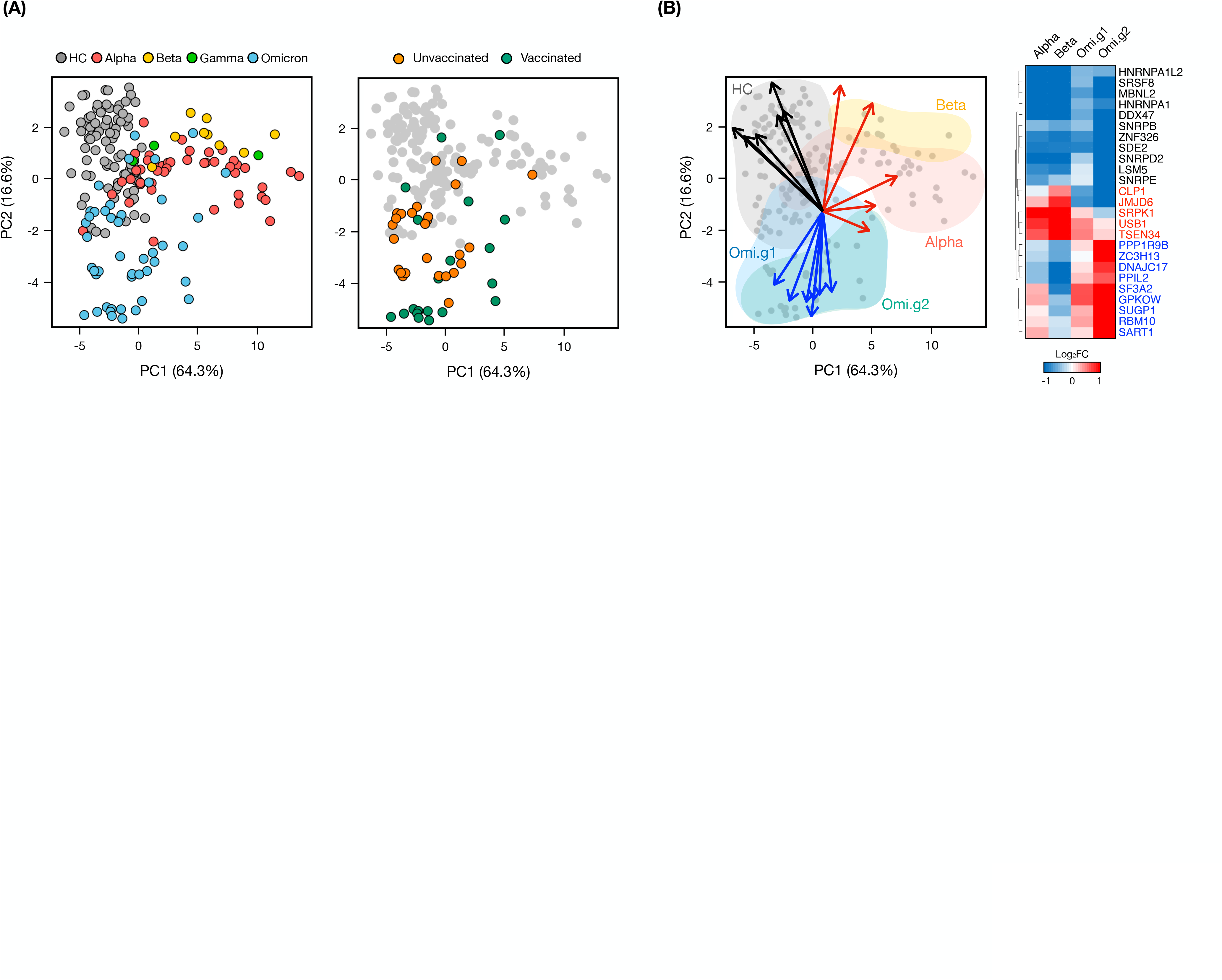
The differentially regulated splicing machinery genes among variants. **(A)** Principal component analysis of 304 genes related to RNA splicing across 190 samples. Each sample are marked HC and COVID-19 patients (left panel) and Omicron patients with vaccination status (right panel). **(B)** Biplot results to identify representative genes with high contributions in each group. (The original biplot results are shown in Fig. S3) Blue arrows represent genes representing the Omicron patient group, red indicates genes representing the Alpha and Beta infection patient groups, and black represents genes representing the HC group. Each gene marked with an arrow on the Biplot is displayed as a heatmap, with values indicating log2 fold change of gene expression levels against HC. Hierarchical clustering was performed with Euclidean distance matrix of fold changes.

## Discussion

In this study, we probed the alternative splicing (AS) landscape in patients infected with various SARS-CoV-2 variants, aiming to illuminate potential underlying mechanisms contributing to the variance in disease manifestations and progression. Our exploratory analysis revealed a significant number of differential alternative splicing events (DASEs), notably in individuals infected with the Omicron variant. This observation necessitates an in-depth investigation into its broader impact on the host’s cellular and molecular responses. Particularly, the observed down-regulation of mRNA splicing machinery genes in COVID-19 patients and their variant-specific modulation signify the critical role of alternative splicing mechanisms in the disease’s progression.

Recent studies have shown that infection by numerous viruses affect to AS landscape of host-cells^34–42^. The modulatory activities of viral products on cellular AS, as reflected through the inhibition of host-cell splicing factor kinases^43–45^ and interference with splicing factors^46,47^ and spliceosome^48–51^, manifest a complex battle between the virus attempting to subvert host defenses and the cell striving to counteract these invasions. This complex interplay prolongs the host-pathogen arms race, with AS serving as a key battleground where success can profoundly affect the infection’s clinical course. The restoration of regular AS patterns may be used to stop viral spread and lessen the severity of the disease if these modulations are understood at a granular level, opening new treatment possibilities.

The re-analysis of 190 RNA-seq datasets, encompassing an average depth of 212 million reads, yielded a rich repository of deep sequencing data. This data depth accords a robust framework to navigate through the complex transcriptional landscape, enabling the detection of genes with low expression levels, and unraveling rare alternative splicing events. Building upon recent studies^13,15,16,49,52^ that highlighted the potential targeting of the mRNA splicing machinery by viruses and reported a dysregulation of alternative splicing in COVID-19 patients, our transcriptomic analysis including healthy controls and patients reinforced the global dynamic alterations of alternative splicing in COVID-19 patients. A notable extent of these changes was observed in those infected with the Omicron variant.

Previous studies have underscored the up-regulation of *TLR4* in severe COVID-19 patients, implicating it in augmented *ACE2* expression and ensuing hyperinflammation^53,54^. Among the alternative splicing forms of *TLR4*, the inclusion of exon 2 or exon 3 has been reported to be elevated in LPS treated cells, mirroring our observation of increased inclusion of these exons in COVID-19 patients^26,27^. Extending the molecular narrative, transcriptome analyses on hospitalized patients infected with Alpha^21^ or Beta^20^ variants revealed a pronounced activation of interferon pathway genes with a spotlight on the JAK/STAT pathway. Concordantly, our analysis unearthed novel DASEs in JAK/STAT genes, hinting at a potential aberrant regulation of the JAK/STAT pathway in COVID-19 patients^55,56^. However, the functional ramifications of these DASEs remain to be elucidated.

Wang et al. reported an increased exclusion form of exon 7 in *CD74* and *LRRFIP1* in the lung tissues of severe COVID-19 patients, alongside a significant down-regulation of six spliceosome component proteins^15^. Our findings corroborated the increased exclusion of *CD74* exon 7 in patients, although we couldn’t confirm this for *LRRFIP1* (Fig. S2). Moreover, our findings point towards a general depression of splicing machinery genes, suggesting a complex interplay far beyond the direct interactions between viral proteins and spliceosome component proteins, potentially disrupting cellular functions and innate immunity^12,35–37^. We also found that the variant- and vaccination-specific gene expression profiles of genes which are member of RNA splicing machinery, suggesting that global and variant specific regulation of AS is highly associated with transcriptional alterations of RNA binding proteins^17,57,58^. This further suggests that the observed splicing gene depression might underlie the abnormal alternative splicing seen in COVID-19 patients.

Numerous genes, including those linked to innate immune activation and strong cytokine activity, have seen an increase in transcription in individuals with severe COVID-19 patients^59–61^. This hyper-activation of transcription might encourage the production of cellular AS transcripts that were not intended^29^. Our study reveals striking alterations in gene expression and AS profiles of numerous immune and cytokine-related genes in COVID-19 patients. Although we observed that AS rates are not correlated with transcription rates, functional studies are needed to know whether AS isoforms increased in patients are functional or transcriptional noise.

Omicron infections generally manifest more moderate symptoms compared to other variants^62^, and vaccinated patients exhibit a significantly blunted interferon response when compared to unvaccinated Omicron infected outpatients and unvaccinated Alpha infected hospitalized patients^63^. On the gene expression front, Omicron showed a gene expression profile like healthy controls compared to Alpha or Beta (Fig. S1), yet alternative splicing exhibited a dynamic alteration, especially pronounced in the vaccinated group. Additionally, although reported in the context of dengue virus vaccination^64^, changes in alternative splicing following vaccination have been documented, suggesting a potential modulation in AS post COVID-19 vaccination. Our observations of T and B cell receptor signaling pathway genes AS changes according to vaccination status hint at the possibility of altered immune cell populations or functionalities.

We reported 25 members of RNA splicing machinery that represent and are specifically expressed in each patient groups. Noteworthy among these is the heterogeneous nuclear ribonucleoprotein A1 (HNRNPA1), known to interact with the 3’-UTR of viruses, orchestrating transcription and replication processes^65–67^. Across all variant groups, *HNRNPA1* exhibited a common downregulation trend. Intriguingly, recent elucidations have spotlighted HNRNPA1 as a hub protein with substantial functional linkages to the human SARS-CoV-2 genome^68^. Furthermore, recent explorations employing various network pharmacology methods have accentuated a close nexus between COVID-19 and serine/arginine-rich splicing factor protein kinase-1 (SRPK1), a gene elucidated to be intimately involved with SARS-CoV-2 replication through the phosphorylation of the N protein^69,70^. Interestingly, our analysis unveiled a pronounced expression of SPRK1 predominantly within the Alpha and Beta infected groups, shedding light on the possible variant-specific molecular dialogues orchestrated by the virus. Furthermore, we delved into the expression patterns of zinc finger CCCH-type containing 13 (ZC3H13), which has been reported to be closely associated with N^6^-methyladenosine (m^6^A), a prevalent epigenetic modification that regulates splicing efficiency^71^ and found in the viral RNA genomes of various viruses^72^. The function of m^6^A in these viral genomes has underscored the intricacies of host-virus interactions at the epigenetic level. Interestingly, a study reported that a lower expression of *ZC3H13* in COVID-19 compared to a higher expression in non-COVID-19 infections^73^. Our analysis unveiled that *ZC3H13* was specifically overexpressed in the Omicron-infected group, particularly within the vaccinated cohort, suggesting a possible interplay between epigenetic modifications and the host’s response to different SARS-CoV-2 variants post-vaccination. Through this exploration, we have laid down a significant marker, directing future research endeavors towards a deeper understanding of the host-virus interactions.

In sum, our investigation unfurls a landscape elucidating the interplay among variant-, vaccination-specific transcriptional changes including gene expression and alternative splicing regulation in the context of SARS-CoV-2 infections. As we delved into the complex realm of alternative splicing, our analysis uncovered alterations that could significantly impact host immune responses, hinting at a critical layer of host-virus interaction that warrants further exploration. Through this comprehensive analysis, we aim to provide a robust framework for understanding how the interplay of viral genetic diversity, host transcriptomic modulation, and vaccination status contribute to the COVID-19 disease spectrum, thereby fostering a more informed foundation for future research and clinical interventions in COVID-19.

## Acknowledgments

This work was supported by the Intramural Research Programs (IRPs) of the National Institute of Diabetes and Digestive and Kidney Diseases, USA.

## Author Contributions

S.-G.L., P.A.F., L.H. and H.K.L designed the study. S.-G.L. analyzed RNA-seq data and administrated the project. S.-G.L., P.A.F., L.H. and H.K.L analyzed data. L.H. and H.K.L. supervised the project. S.-G.L., P.A.F., L.H. and H.K.L wrote the paper. All authors read and approved the manuscript.

## STAR★Methods

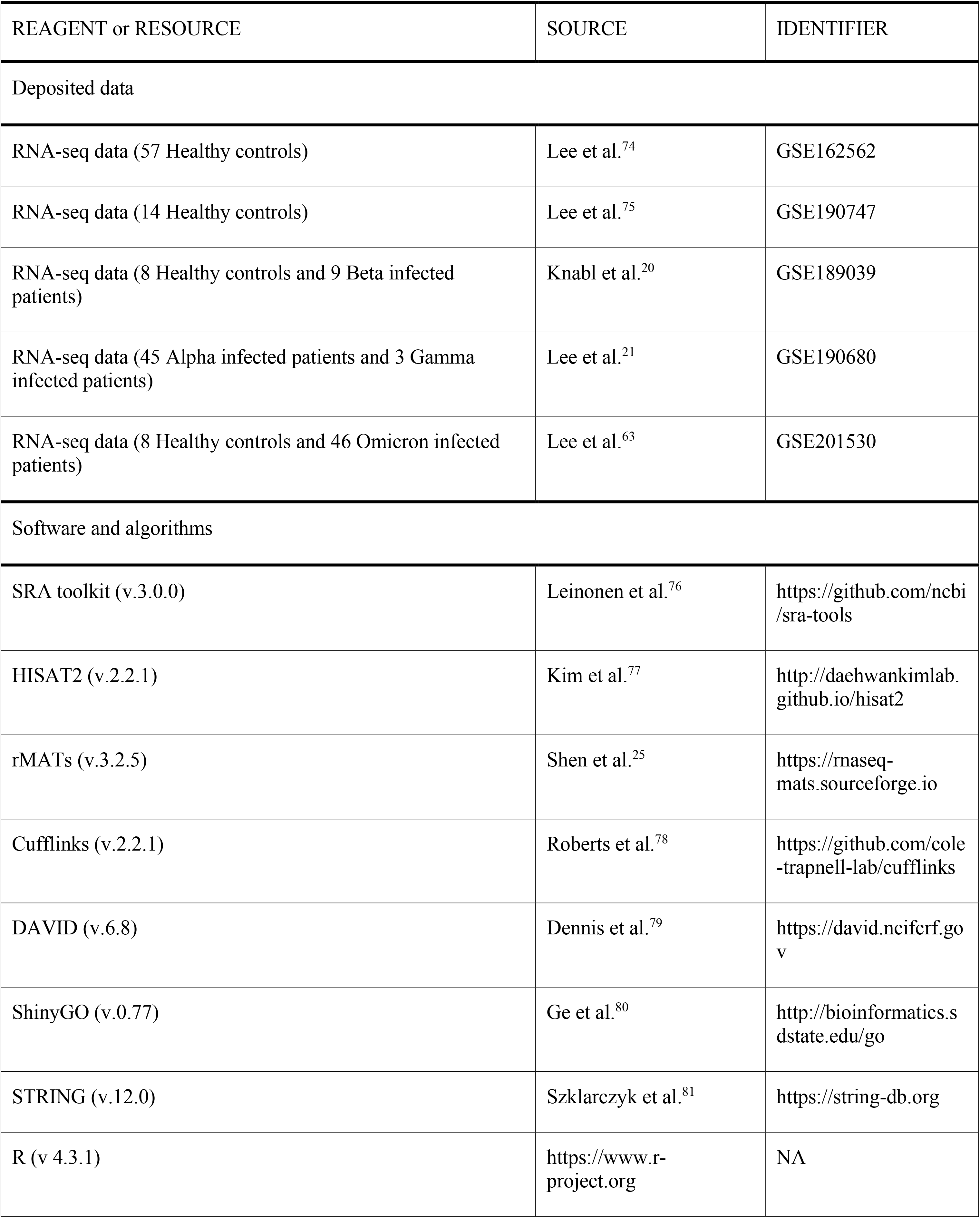
Key resources table.

## Resource availability

### Lead contact

Further information and requests for resources should be directed to the lead contact, Hye Kyung Lee.

### Materials availability

This study did not generate new unique reagents.

## Method details

### Data collection

We obtained whole blood RNA-seq data from COVID-19 patients infected with four different variants (Alpha, Beta, Gamma, and Omicron), as well as from a group of healthy controls, using data from five Gene Expression Omnibus (GEO) datasets (see Table S1 for details)^20–22,63,74,75^. The healthy control group comprised 87 samples, while the COVID-19 patient group consisted of 103 samples. To focus on early-stage infection, we exclusively collected data from patients who had been infected for less than one week. All raw sequences were downloaded and converted to FASTQ by SRA toolkit (v.3.0.0)^76^.

### Alternative splicing and transcriptome analyses

The raw reads were initially subjected to preprocessing, involving the removal of poor-quality 3’ ends, utilizing the *trimFastq.py* script, which is an integral component of rMATs (v.3.2.5)^25^. Subsequently, the resulting cleaned reads were then mapped to the human genome using HISAT2 (v.2.2.1)^77^ with default parameter settings. Furthermore, an HISAT2 genome index was constructed as part of the preparation for this mapping step. To identify alternative splicing events, we ran rMATs with default parameters and human gene annotations. We utilized a widely recognized exon-based ratio metric known as the percent spliced in index (PSI) ratio to quantify alternative splicing events. The PSI ratio is calculated as follows:

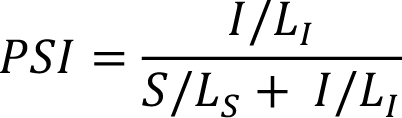

where *S* and *I* represents the number of reads mapped to the junction supporting the skipping and inclusion form, respectively. *L* signifies the effective length, which is used for normalization.

To identify differential alternative splicing events (DASEs), we conducted a comparison of exon inclusion levels between healthy control and COVID-19 patients for each alternative splicing event. DASEs were defined using the following criteria: |PSI differences| > 0.1 and corrected *P*-value < 0.05. For the statistical analysis, we employed the Wilcoxon rank-sum test, and obtained corrected *p*-value by Bonferroni correction.

To quantify gene expression levels, we employed Cufflinks (v.2.2.1)^78^ to assemble genome-aligned reads into transcripts. The relative abundances of these transcripts were estimated based on the read count support for each transcript. Unless otherwise stated, we utilized fragments per kilobase per million reads mapped (FPKM) as the unit of gene expression levels. Differentially expressed genes (DEGs) were defined as genes with a significant expression difference between groups, characterized by a corrected p-value lower than 0.05 in group-to-group comparisons. The human reference genome sequence and annotation files were acquired from the UCSC genome browser (https://genome.ucsc.edu) under version hg38. Additionally, we excluded uncharacterized and alternative chromosomes and their associated genes from our analysis.

### Gene set enrichment and over-representation analysis

We performed over-representation analysis for pathway enrichment of DASEs and DEGs using DAVID (v.6.8)^79^ (https://david.ncifcrf.gov) and ShinyGO (v.0.77)^80^ (http://bioinformatics.sdstate.edu/go). In this analysis, we selected significant categories and pathways with a false discovery rate (FDR) less than 0.05 for further investigation. The gene set of gene ontology (GO) terms and pathways were obtained Gene Ontology resources (https://geneontology.org) and Kyoto Encyclopedia of Genes and Genomes (KEGG) pathway database (https://www.genome.jp/kegg/pathway).

### SARS-CoV-2 susceptible genes

SARS-CoV-2 susceptible genes were collected form PanelApp^82^, a resource that provides curated information about genes and their associations with specific diseases and conditions, specifically utilizing the panel of COVID-19 research (v.1.136) (https://panelapp.genomicsengland.co.uk/panels/111).

### Protein-protein interaction (PPI) network analysis

We construct a PPI network from DEGs of COVID-19 patients, specifically enriched in RNA splicing term (GO:0008380) by STRING database (v.12.0)^81^ (https://string-db.org). We employed stringent parameters, requiring interactions with the highest confidence (minimum required interaction score > 0.9) and hiding disconnected nodes in the network.

## Quantification and statistical analysis

### Data analysis

Principal component analysis (PCA) was performed using the exon inclusion levels of all alternative splicing events by *prcomp* function from R *stat* package with the first two principal components. To cluster the samples based on alternative splicing profiles, we conducted *k*-means clustering, which is one of the unsupervised clustering methods. To determine the optimal number of clusters (*k*), we performed multiple analyses by setting k from 2 to 5. Through this iterative process, we examined the saturation points of the total within sum of squares (SS) values and identified that the optimal value for *k* is 2. To assess the degree of gene contributions to principal components, we utilized *biplots* from the R *stat* package. The *p* values from comparing distributions were obtained by Wilcoxon rank-sum test. All *p* values were adjusted by Bonferroni correction.

## Data and code availability

- Differentially alternative spliced events (DASEs) data from this study is available on our GitHub repository: https://github.com/tjdrnjsqpf/COVID19_AS.
- This paper does not report original code.
- Any additional information required to reanalyze the data reported in this paper is available from the lead contact upon request.

## Supplementary information

**Document S1. Figures S1-S3**

**Table S1. RNA-seq data statistics**

## References

1. Feng, Y., Ling, Y., Bai, T., Xie, Y., Huang, J., Li, J., Xiong, W., Yang, D., Chen, R., Lu, F., et al. (2020). COVID-19 with Different Severities: A Multicenter Study of Clinical Features. Am J Respir Crit Care Med 201, 1380–1388. 10.1164/rccm.202002-0445OC.

2. Liang, W.H., Guan, W.J., Li, C.C., Li, Y.M., Liang, H.R., Zhao, Y., Liu, X.Q., Sang, L., Chen, R.C., Tang, C.L., et al. (2020). Clinical characteristics and outcomes of hospitalised patients with COVID-19 treated in Hubei (epicentre) and outside Hubei (non-epicentre): a nationwide analysis of China. Eur Respir J 55. 10.1183/13993003.00562-2020.

3. Zhou, F., Yu, T., Du, R., Fan, G., Liu, Y., Liu, Z., Xiang, J., Wang, Y., Song, B., Gu, X., et al. (2020). Clinical course and risk factors for mortality of adult inpatients with COVID-19 in Wuhan, China: a retrospective cohort study. Lancet 395, 1054–1062. 10.1016/S0140-6736(20)30566-3.

4. Chen, G., Wu, D., Guo, W., Cao, Y., Huang, D., Wang, H., Wang, T., Zhang, X., Chen, H., Yu, H., et al. (2020). Clinical and immunological features of severe and moderate coronavirus disease 2019. J Clin Invest 130, 2620–2629. 10.1172/JCI137244.

5. Pascarella, G., Strumia, A., Piliego, C., Bruno, F., Del Buono, R., Costa, F., Scarlata, S., and Agro, F.E. (2020). COVID-19 diagnosis and management: a comprehensive review. J Intern Med 288, 192–206. 10.1111/joim.13091.

6. Bojkova, D., Klann, K., Koch, B., Widera, M., Krause, D., Ciesek, S., Cinatl, J., and Munch, C. (2020). Proteomics of SARS-CoV-2-infected host cells reveals therapy targets. Nature 583, 469–472. 10.1038/s41586-020-2332-7.

7. Li, C.X., Gao, J., Zhang, Z., Chen, L., Li, X., Zhou, M., and Wheelock, A.M. (2022). Multiomics integration-based molecular characterizations of COVID-19. Brief Bioinform 23. 10.1093/bib/bbab485.

8. Wu, M., Chen, Y., Xia, H., Wang, C., Tan, C.Y., Cai, X., Liu, Y., Ji, F., Xiong, P., Liu, R., et al. (2020). Transcriptional and proteomic insights into the host response in fatal COVID-19 cases. Proc Natl Acad Sci U S A 117, 28336–28343. 10.1073/pnas.2018030117.

9. Kim, E., Magen, A., and Ast, G. (2007). Different levels of alternative splicing among eukaryotes. Nucleic Acids Res 35, 125–131. 10.1093/nar/gkl924.

10. Nilsen, T.W., and Graveley, B.R. (2010). Expansion of the eukaryotic proteome by alternative splicing. Nature 463, 457–463. 10.1038/nature08909.

11. Mukherjee, S.B., Mukherjee, S., Detroja, R., and Frenkel-Morgenstern, M. (2023). The landscape of differential splicing and transcript alternations in severe COVID-19 infection. FEBS J 290, 3128–3144. 10.1111/febs.16723.

12. Ashraf, U., Benoit-Pilven, C., Lacroix, V., Navratil, V., and Naffakh, N. (2019). Advances in Analyzing Virus-Induced Alterations of Host Cell Splicing. Trends Microbiol 27, 268–281. 10.1016/j.tim.2018.11.004.

13. Karlebach, G., Aronow, B., Baylin, S.B., Butler, D., Foox, J., Levy, S., Meydan, C., Mozsary, C., Saravia-Butler, A.M., Taylor, D.M., et al. (2022). Betacoronavirus-specific alternate splicing. Genomics 114, 110270. 10.1016/j.ygeno.2022.110270.

14. Srivastava, R., Daulatabad, S.V., Srivastava, M., and Janga, S.C. (2020). Role of SARS-CoV-2 in altering the RNA binding protein and miRNA directed post-transcriptional regulatory networks in humans. bioRxiv. 10.1101/2020.07.06.190348.

15. Wang, C., Chen, L., Chen, Y., Jia, W., Cai, X., Liu, Y., Ji, F., Xiong, P., Liang, A., Liu, R., et al. (2022). Abnormal global alternative RNA splicing in COVID-19 patients. PLoS Genet 18, e1010137. 10.1371/journal.pgen.1010137.

16. Banerjee, A.K., Blanco, M.R., Bruce, E.A., Honson, D.D., Chen, L.M., Chow, A., Bhat, P., Ollikainen, N., Quinodoz, S.A., Loney, C., et al. (2020). SARS-CoV-2 Disrupts Splicing, Translation, and Protein Trafficking to Suppress Host Defenses. Cell 183, 1325–1339 e1321. 10.1016/j.cell.2020.10.004.

17. Huang, R., Chen, W., Zhao, X., Ma, Y., Zhou, Q., Chen, J., Zhang, M., Zhao, D., Hou, Y., He, C., and Wu, Y. (2023). Genome-wide characterization of alternative splicing in blood cells of COVID-19 and respiratory infections of relevance. Virol Sin 38, 309–312. 10.1016/j.virs.2023.01.007.

18. Huffman, J., Butler-Laporte, G., Khan, A., Drivas, T.G., Peloso, G.M., Nakanishi, T., Verma, A., Kiryluk, K., Richards, J.B., and Zeberg, H. (2021). Alternative splicing of OAS1 alters the risk for severe COVID-19. medRxiv. 10.1101/2021.03.20.21254005.

19. Knabl, L., Lee, H.K., Wieser, M., Mur, A., Zabemigg, A., Knabl, L., Sr., Rauch, S., Bock, M., Schumacher, J., Kaiser, N., Furth, P., and Hennighausen, L. (2021). Impact of BNT162b First Vaccination on the Immune Transcriptome of Elderly Patients Infected with the B.1.351 SARS-CoV-2 Variant. Res Sq. 10.21203/rs.3.rs-509143/v1.

20. Knabl, L., Lee, H.K., Wieser, M., Mur, A., Zabernigg, A., Knabl, L., Sr., Rauch, S., Bock, M., Schumacher, J., Kaiser, N., Furth, P.A., and Hennighausen, L. (2022). BNT162b2 vaccination enhances interferon-JAK-STAT-regulated antiviral programs in COVID-19 patients infected with the SARS-CoV-2 Beta variant. Commun Med (Lond) 2. 10.1038/s43856-022-00083-x.

21. Lee, H.K., Knabl, L., Knabl, L., Sr., Wieser, M., Mur, A., Zabernigg, A., Schumacher, J., Kapferer, S., Kaiser, N., Furth, P.A., and Hennighausen, L. (2022). Immune transcriptome analysis of COVID-19 patients infected with SARS-CoV-2 variants carrying the E484K escape mutation identifies a distinct gene module. Sci Rep 12, 2784. 10.1038/s41598-022-06752-0.

22. Lee, H.K., Knabl, L., Walter, M., Furth, P.A., and Hennighausen, L. (2022). Limited cross-variant immune response from SARS-CoV-2 Omicron BA.2 in naive but not previously infected outpatients. iScience 25, 105369. 10.1016/j.isci.2022.105369.

23. Mistry, P., Barmania, F., Mellet, J., Peta, K., Strydom, A., Viljoen, I.M., James, W., Gordon, S., and Pepper, M.S. (2021). SARS-CoV-2 Variants, Vaccines, and Host Immunity. Front Immunol 12, 809244. 10.3389/fimmu.2021.809244.

24. Wang, H., Liu, C., Xie, X., Niu, M., Wang, Y., Cheng, X., Zhang, B., Zhang, D., Liu, M., Sun, R., et al. (2023). Multi-omics blood atlas reveals unique features of immune and platelet responses to SARS-CoV-2 Omicron breakthrough infection. Immunity 56, 1410–1428 e1418. 10.1016/j.immuni.2023.05.007.

25. Shen, S., Park, J.W., Lu, Z.X., Lin, L., Henry, M.D., Wu, Y.N., Zhou, Q., and Xing, Y. (2014). rMATS: robust and flexible detection of differential alternative splicing from replicate RNA-Seq data. Proc Natl Acad Sci U S A 111, E5593–5601. 10.1073/pnas.1419161111.

26. Iwami, K.I., Matsuguchi, T., Masuda, A., Kikuchi, T., Musikacharoen, T., and Yoshikai, Y. (2000). Cutting edge: naturally occurring soluble form of mouse Toll-like receptor 4 inhibits lipopolysaccharide signaling. J Immunol 165, 6682–6686. 10.4049/jimmunol.165.12.6682.

27. Jaresova, I., Rozkova, D., Spisek, R., Janda, A., Brazova, J., and Sediva, A. (2007). Kinetics of Toll-like receptor-4 splice variants expression in lipopolysaccharide-stimulated antigen presenting cells of healthy donors and patients with cystic fibrosis. Microbes Infect 9, 1359–1367. 10.1016/j.micinf.2007.06.009.

28. Ravid, J.D., Leiva, O., and Chitalia, V.C. (2022). Janus Kinase Signaling Pathway and Its Role in COVID-19 Inflammatory, Vascular, and Thrombotic Manifestations. Cells 11. 10.3390/cells11020306.

29. Aslanzadeh, V., Huang, Y., Sanguinetti, G., and Beggs, J.D. (2018). Transcription rate strongly affects splicing fidelity and cotranscriptionality in budding yeast. Genome Res 28, 203–213. 10.1101/gr.225615.117.

30. Nuevo-Tapioles, C., and Philips, M.R. (2022). The role of KRAS splice variants in cancer biology. Front Cell Dev Biol 10, 1033348. 10.3389/fcell.2022.1033348.

31. Riffo-Campos, A.L., Gimeno-Valiente, F., Rodriguez, F.M., Cervantes, A., Lopez-Rodas, G., Franco, L., and Castillo, J. (2018). Role of epigenetic factors in the selection of the alternative splicing isoforms of human KRAS in colorectal cancer cell lines. Oncotarget 9, 20578–20589. 10.18632/oncotarget.25016.

32. Voice, J.K., Klemke, R.L., Le, A., and Jackson, J.H. (1999). Four human ras homologs differ in their abilities to activate Raf-1, induce transformation, and stimulate cell motility. J Biol Chem 274, 17164–17170. 10.1074/jbc.274.24.17164.

33. Vihma, H., Pruunsild, P., and Timmusk, T. (2008). Alternative splicing and expression of human and mouse NFAT genes. Genomics 92, 279–291. 10.1016/j.ygeno.2008.06.011.

34. Batra, R., Stark, T.J., Clark, A.E., Belzile, J.P., Wheeler, E.C., Yee, B.A., Huang, H., Gelboin-Burkhart, C., Huelga, S.C., Aigner, S., et al. (2016). RNA-binding protein CPEB1 remodels host and viral RNA landscapes. Nat Struct Mol Biol 23, 1101–1110. 10.1038/nsmb.3310.

35. Boudreault, S., Armero, V.E.S., Scott, M.S., Perreault, J.P., and Bisaillon, M. (2019). The Epstein-Barr virus EBNA1 protein modulates the alternative splicing of cellular genes. Virol J 16, 29. 10.1186/s12985-019-1137-5.

36. Boudreault, S., Martenon-Brodeur, C., Caron, M., Garant, J.M., Tremblay, M.P., Armero, V.E., Durand, M., Lapointe, E., Thibault, P., Tremblay-Letourneau, M., et al. (2016). Global Profiling of the Cellular Alternative RNA Splicing Landscape during Virus-Host Interactions. PLoS One 11, e0161914. 10.1371/journal.pone.0161914.

37. Boudreault, S., Roy, P., Lemay, G., and Bisaillon, M. (2019). Viral modulation of cellular RNA alternative splicing: A new key player in virus-host interactions? Wiley Interdiscip Rev RNA 10, e1543. 10.1002/wrna.1543.

38. Hu, B., Huo, Y., Yang, L., Chen, G., Luo, M., Yang, J., and Zhou, J. (2017). ZIKV infection effects changes in gene splicing, isoform composition and lncRNA expression in human neural progenitor cells. Virol J 14, 217. 10.1186/s12985-017-0882-6.

39. Thenoz, M., Vernin, C., Mortada, H., Karam, M., Pinatel, C., Gessain, A., Webb, T.R., Auboeuf, D., Wattel, E., and Mortreux, F. (2014). HTLV-1-infected CD4+ T-cells display alternative exon usages that culminate in adult T-cell leukemia. Retrovirology 11, 119. 10.1186/s12977-014-0119-3.

40. Thompson, M.G., Dittmar, M., Mallory, M.J., Bhat, P., Ferretti, M.B., Fontoura, B.M., Cherry, S., and Lynch, K.W. (2020). Viral-induced alternative splicing of host genes promotes influenza replication. Elife 9. 10.7554/eLife.55500.

41. Thompson, M.G., Munoz-Moreno, R., Bhat, P., Roytenberg, R., Lindberg, J., Gazzara, M.R., Mallory, M.J., Zhang, K., Garcia-Sastre, A., Fontoura, B.M.A., and Lynch, K.W. (2018). Co-regulatory activity of hnRNP K and NS1-BP in influenza and human mRNA splicing. Nat Commun 9, 2407. 10.1038/s41467-018-04779-4.

42. Xu, J., Fang, Y., Qin, J., Chen, X., Liang, X., Xie, X., and Lu, W. (2016). A transcriptomic landscape of human papillomavirus 16 E6-regulated gene expression and splicing events. FEBS Lett 590, 4594–4605. 10.1002/1873-3468.12486.

43. Lindberg, A., and Kreivi, J.P. (2002). Splicing inhibition at the level of spliceosome assembly in the presence of herpes simplex virus protein ICP27. Virology 294, 189–198. 10.1006/viro.2001.1301.

44. Phelan, A., Carmo-Fonseca, M., McLaughlan, J., Lamond, A.I., and Clements, J.B. (1993). A herpes simplex virus type 1 immediate-early gene product, IE63, regulates small nuclear ribonucleoprotein distribution. Proc Natl Acad Sci U S A 90, 9056–9060. 10.1073/pnas.90.19.9056.

45. Sciabica, K.S., Dai, Q.J., and Sandri-Goldin, R.M. (2003). ICP27 interacts with SRPK1 to mediate HSV splicing inhibition by altering SR protein phosphorylation. EMBO J 22, 1608–1619. 10.1093/emboj/cdg166.

46. Verma, D., Bais, S., Gaillard, M., and Swaminathan, S. (2010). Epstein-Barr Virus SM protein utilizes cellular splicing factor SRp20 to mediate alternative splicing. J Virol 84, 11781–11789. 10.1128/JVI.01359-10.

47. Verma, D., and Swaminathan, S. (2008). Epstein-Barr virus SM protein functions as an alternative splicing factor. J Virol 82, 7180–7188. 10.1128/JVI.00344-08.

48. Chiba, S., Hill-Batorski, L., Neumann, G., and Kawaoka, Y. (2018). The Cellular DExD/H-Box RNA Helicase UAP56 Co-localizes With the Influenza A Virus NS1 Protein. Front Microbiol 9, 2192. 10.3389/fmicb.2018.02192.

49. De Maio, F.A., Risso, G., Iglesias, N.G., Shah, P., Pozzi, B., Gebhard, L.G., Mammi, P., Mancini, E., Yanovsky, M.J., Andino, R., et al. (2016). The Dengue Virus NS5 Protein Intrudes in the Cellular Spliceosome and Modulates Splicing. PLoS Pathog 12, e1005841. 10.1371/journal.ppat.1005841.

50. Fabozzi, G., Oler, A.J., Liu, P., Chen, Y., Mindaye, S., Dolan, M.A., Kenney, H., Gucek, M., Zhu, J., Rabin, R.L., and Subbarao, K. (2018). Strand-Specific Dual RNA Sequencing of Bronchial Epithelial Cells Infected with Influenza A/H3N2 Viruses Reveals Splicing of Gene Segment 6 and Novel Host-Virus Interactions. J Virol 92. 10.1128/JVI.00518-18.

51. Hashizume, C., Kuramitsu, M., Zhang, X., Kurosawa, T., Kamata, M., and Aida, Y. (2007). Human immunodeficiency virus type 1 Vpr interacts with spliceosomal protein SAP145 to mediate cellular pre-mRNA splicing inhibition. Microbes Infect 9, 490–497. 10.1016/j.micinf.2007.01.013.

52. An, S., Li, Y., Lin, Y., Chu, J., Su, J., Chen, Q., Wang, H., Pan, P., Zheng, R., Li, J., et al. (2021). Genome-Wide Profiling Reveals Alternative Polyadenylation of Innate Immune-Related mRNA in Patients With COVID-19. Front Immunol 12, 756288. 10.3389/fimmu.2021.756288.

53. Aboudounya, M.M., and Heads, R.J. (2021). COVID-19 and Toll-Like Receptor 4 (TLR4): SARS-CoV-2 May Bind and Activate TLR4 to Increase ACE2 Expression, Facilitating Entry and Causing Hyperinflammation. Mediators Inflamm 2021, 8874339. 10.1155/2021/8874339.

54. Sohn, K.M., Lee, S.G., Kim, H.J., Cheon, S., Jeong, H., Lee, J., Kim, I.S., Silwal, P., Kim, Y.J., Paik, S., et al. (2020). COVID-19 Patients Upregulate Toll-like Receptor 4-mediated Inflammatory Signaling That Mimics Bacterial Sepsis. J Korean Med Sci 35, e343. 10.3346/jkms.2020.35.e343.

55. Luo, J., Lu, S., Yu, M., Zhu, L., Zhu, C., Li, C., Fang, J., Zhu, X., and Wang, X. (2021). The potential involvement of JAK-STAT signaling pathway in the COVID-19 infection assisted by ACE2. Gene 768, 145325. 10.1016/j.gene.2020.145325.

56. Matsuyama, T., Kubli, S.P., Yoshinaga, S.K., Pfeffer, K., and Mak, T.W. (2020). An aberrant STAT pathway is central to COVID-19. Cell Death Differ 27, 3209–3225. 10.1038/s41418-020-00633-7.

57. Fu, X.D., and Ares, M., Jr. (2014). Context-dependent control of alternative splicing by RNA-binding proteins. Nat Rev Genet 15, 689–701. 10.1038/nrg3778.

58. Ule, J., and Blencowe, B.J. (2019). Alternative Splicing Regulatory Networks: Functions, Mechanisms, and Evolution. Mol Cell 76, 329–345. 10.1016/j.molcel.2019.09.017.

59. Karki, R., and Kanneganti, T.D. (2022). Innate immunity, cytokine storm, and inflammatory cell death in COVID-19. J Transl Med 20, 542. 10.1186/s12967-022-03767-z.

60. Lowery, S.A., Sariol, A., and Perlman, S. (2021). Innate immune and inflammatory responses to SARS-CoV-2: Implications for COVID-19. Cell Host Microbe 29, 1052–1062. 10.1016/j.chom.2021.05.004.

61. Teijaro, J.R., and Farber, D.L. (2021). COVID-19 vaccines: modes of immune activation and future challenges. Nat Rev Immunol 21, 195–197. 10.1038/s41577-021-00526-x.

62. Petersen, M.S., S, I.K., Eliasen, E.H., Larsen, S., Hansen, J.L., Vest, N., Dahl, M.M., Christiansen, D.H., Moller, L.F., and Kristiansen, M.F. (2022). Clinical characteristics of the Omicron variant - results from a Nationwide Symptoms Survey in the Faroe Islands. Int J Infect Dis 122, 636–643. 10.1016/j.ijid.2022.07.005.

63. Lee, H.K., Knabl, L., Walter, M., Knabl, L., Sr., Dai, Y., Fussl, M., Caf, Y., Jeller, C., Knabl, P., Obermoser, M., et al. (2022). Prior Vaccination Exceeds Prior Infection in Eliciting Innate and Humoral Immune Responses in Omicron Infected Outpatients. Front Immunol 13, 916686. 10.3389/fimmu.2022.916686.

64. Kim, E.Y., Che, Y., Dean, H.J., Lorenzo-Redondo, R., Stewart, M., Keller, C.K., Whorf, D., Mills, D., Dulin, N.N., Kim, T., et al. (2022). Transcriptome-wide changes in gene expression, splicing, and lncRNAs in response to a live attenuated dengue virus vaccine. Cell Rep 38, 110341. 10.1016/j.celrep.2022.110341.

65. Huang, P., and Lai, M.M. (2001). Heterogeneous nuclear ribonucleoprotein a1 binds to the 3’-untranslated region and mediates potential 5’-3’-end cross talks of mouse hepatitis virus RNA. J Virol 75, 5009–5017. 10.1128/JVI.75.11.5009-5017.2001.

66. Li, H.P., Zhang, X., Duncan, R., Comai, L., and Lai, M.M. (1997). Heterogeneous nuclear ribonucleoprotein A1 binds to the transcription-regulatory region of mouse hepatitis virus RNA. Proc Natl Acad Sci U S A 94, 9544–9549. 10.1073/pnas.94.18.9544.

67. Shi, S.T., Huang, P., Li, H.P., and Lai, M.M. (2000). Heterogeneous nuclear ribonucleoprotein A1 regulates RNA synthesis of a cytoplasmic virus. EMBO J 19, 4701–4711. 10.1093/emboj/19.17.4701.

68. Zhou, Y., Hou, Y., Shen, J., Huang, Y., Martin, W., and Cheng, F. (2020). Network-based drug repurposing for novel coronavirus 2019-nCoV/SARS-CoV-2. Cell Discov 6, 14. 10.1038/s41421-020-0153-3.

69. Wang, Z., Zhan, J., and Gao, H. (2022). Computer-aided drug design combined network pharmacology to explore anti-SARS-CoV-2 or anti-inflammatory targets and mechanisms of Qingfei Paidu Decoction for COVID-19. Front Immunol 13, 1015271. 10.3389/fimmu.2022.1015271.

70. Yaron, T.M., Heaton, B.E., Levy, T.M., Johnson, J.L., Jordan, T.X., Cohen, B.M., Kerelsky, A., Lin, T.Y., Liberatore, K.M., Bulaon, D.K., et al. (2020). The FDA-approved drug Alectinib compromises SARS-CoV-2 nucleocapsid phosphorylation and inhibits viral infection in vitro. bioRxiv. 10.1101/2020.08.14.251207.

71. Louloupi, A., Ntini, E., Conrad, T., and Orom, U.A.V. (2018). Transient N-6-Methyladenosine Transcriptome Sequencing Reveals a Regulatory Role of m6A in Splicing Efficiency. Cell Rep 23, 3429–3437. 10.1016/j.celrep.2018.05.077.

72. Wen, J., Lv, R., Ma, H., Shen, H., He, C., Wang, J., Jiao, F., Liu, H., Yang, P., Tan, L., et al. (2018). Zc3h13 Regulates Nuclear RNA m(6)A Methylation and Mouse Embryonic Stem Cell Self-Renewal. Mol Cell 69, 1028–1038 e1026. 10.1016/j.molcel.2018.02.015.

73. Thair, S.A., He, Y.D., Hasin-Brumshtein, Y., Sakaram, S., Pandya, R., Toh, J., Rawling, D., Remmel, M., Coyle, S., Dalekos, G.N., et al. (2021). Transcriptomic similarities and differences in host response between SARS-CoV-2 and other viral infections. iScience 24, 101947. 10.1016/j.isci.2020.101947.

74. Lee, H.K., Knabl, L., Pipperger, L., Volland, A., Furth, P.A., Kang, K., Smith, H.E., Knabl, L., Sr., Bellmann, R., Bernhard, C., et al. (2021). Immune transcriptomes of highly exposed SARS-CoV-2 asymptomatic seropositive versus seronegative individuals from the Ischgl community. Sci Rep 11, 4243. 10.1038/s41598-021-83110-6.

75. Lee, H.K., Knabl, L., Moliva, J.I., Knabl, L., Sr., Werner, A.P., Boyoglu-Barnum, S., Kapferer, S., Pateter, B., Walter, M., Sullivan, N.J., Furth, P.A., and Hennighausen, L. (2022). mRNA vaccination in octogenarians 15 and 20 months after recovery from COVID-19 elicits robust immune and antibody responses that include Omicron. Cell Rep 39, 110680. 10.1016/j.celrep.2022.110680.

76. Leinonen, R., Sugawara, H., Shumway, M., and International Nucleotide Sequence Database, C. (2011). The sequence read archive. Nucleic Acids Res 39, D19–21. 10.1093/nar/gkq1019.

77. Kim, D., Paggi, J.M., Park, C., Bennett, C., and Salzberg, S.L. (2019). Graph-based genome alignment and genotyping with HISAT2 and HISAT-genotype. Nat Biotechnol 37, 907–915. 10.1038/s41587-019-0201-4.

78. Roberts, A., Pimentel, H., Trapnell, C., and Pachter, L. (2011). Identification of novel transcripts in annotated genomes using RNA-Seq. Bioinformatics 27, 2325–2329. 10.1093/bioinformatics/btr355.

79. Dennis, G., Jr., Sherman, B.T., Hosack, D.A., Yang, J., Gao, W., Lane, H.C., and Lempicki, R.A. (2003). DAVID: Database for Annotation, Visualization, and Integrated Discovery. Genome Biol 4, P3.

80. Ge, S.X., Jung, D., and Yao, R. (2020). ShinyGO: a graphical gene-set enrichment tool for animals and plants. Bioinformatics 36, 2628–2629. 10.1093/bioinformatics/btz931.

81. Szklarczyk, D., Gable, A.L., Nastou, K.C., Lyon, D., Kirsch, R., Pyysalo, S., Doncheva, N.T., Legeay, M., Fang, T., Bork, P., Jensen, L.J., and von Mering, C. (2021). The STRING database in 2021: customizable protein-protein networks, and functional characterization of user-uploaded gene/measurement sets. Nucleic Acids Res 49, D605–D612. 10.1093/nar/gkaa1074.

82. Martin, A.R., Williams, E., Foulger, R.E., Leigh, S., Daugherty, L.C., Niblock, O., Leong, I.U.S., Smith, K.R., Gerasimenko, O., Haraldsdottir, E., et al. (2019). PanelApp crowdsources expert knowledge to establish consensus diagnostic gene panels. Nat Genet 51, 1560–1565. 10.1038/s41588-019-0528-2.

