## Supplementary figures for "Variant- and Vaccination-Specific Alternative Splicing Profiles in SARS-CoV-2 Infections"

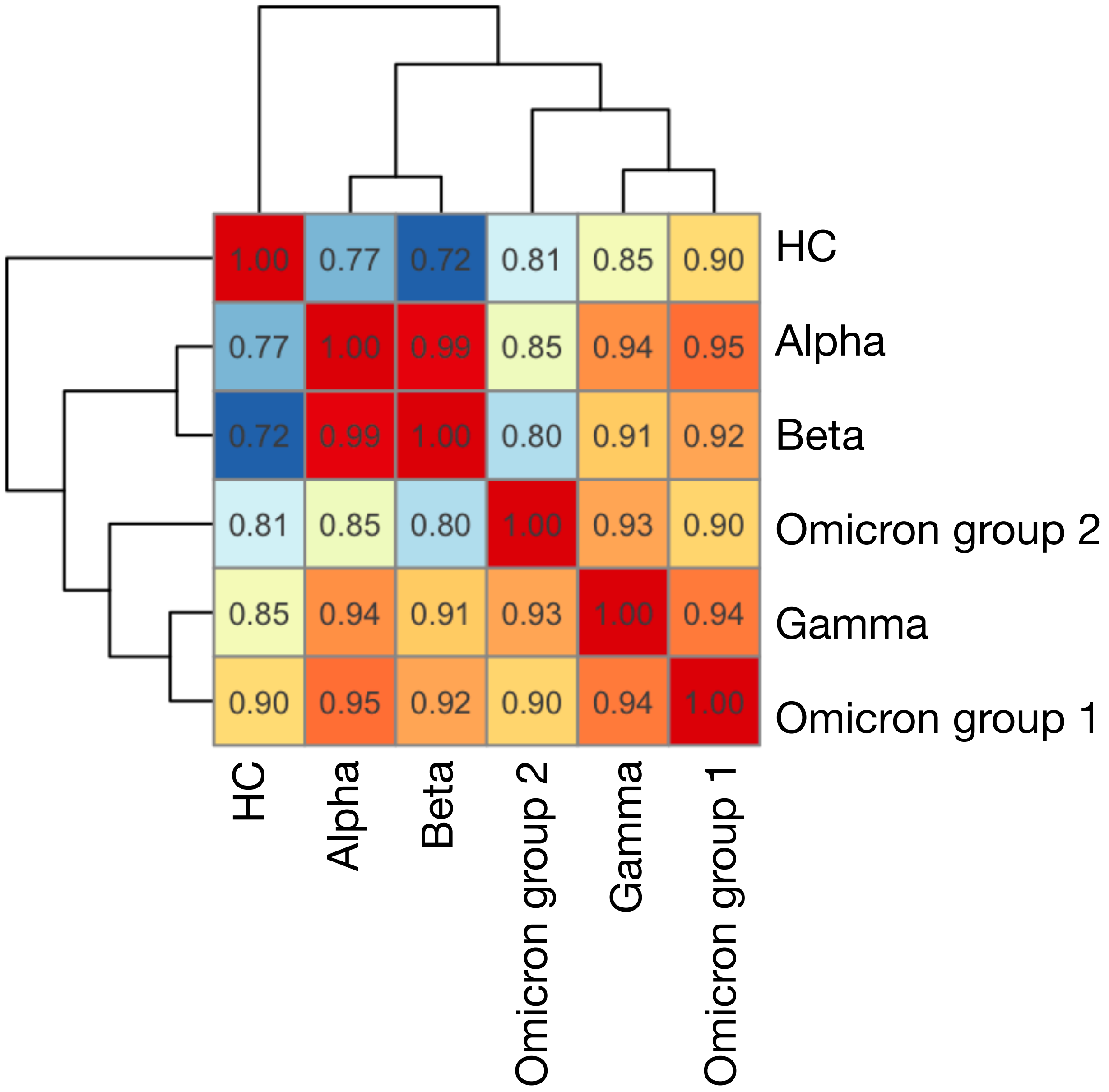


Supplementary figure 1. Gene expression level correlation between groups. Each value indicates Pearson's correlation coefficient.

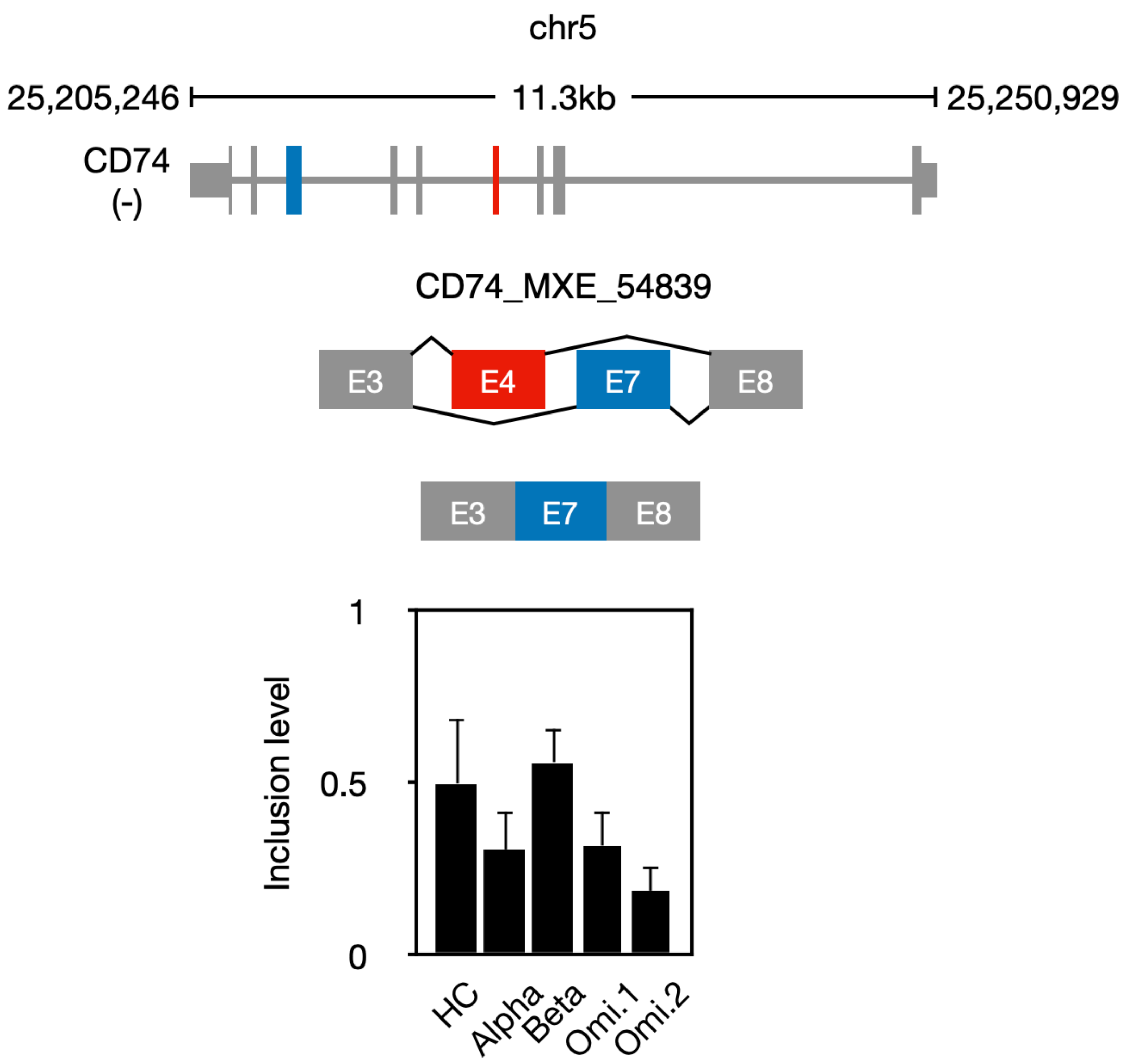


Supplementary figure 2. DASEs of *CD74* of which exon 7 significantly excluded in COVID-19 patients. The error bar indicates standard deviation of the mean.

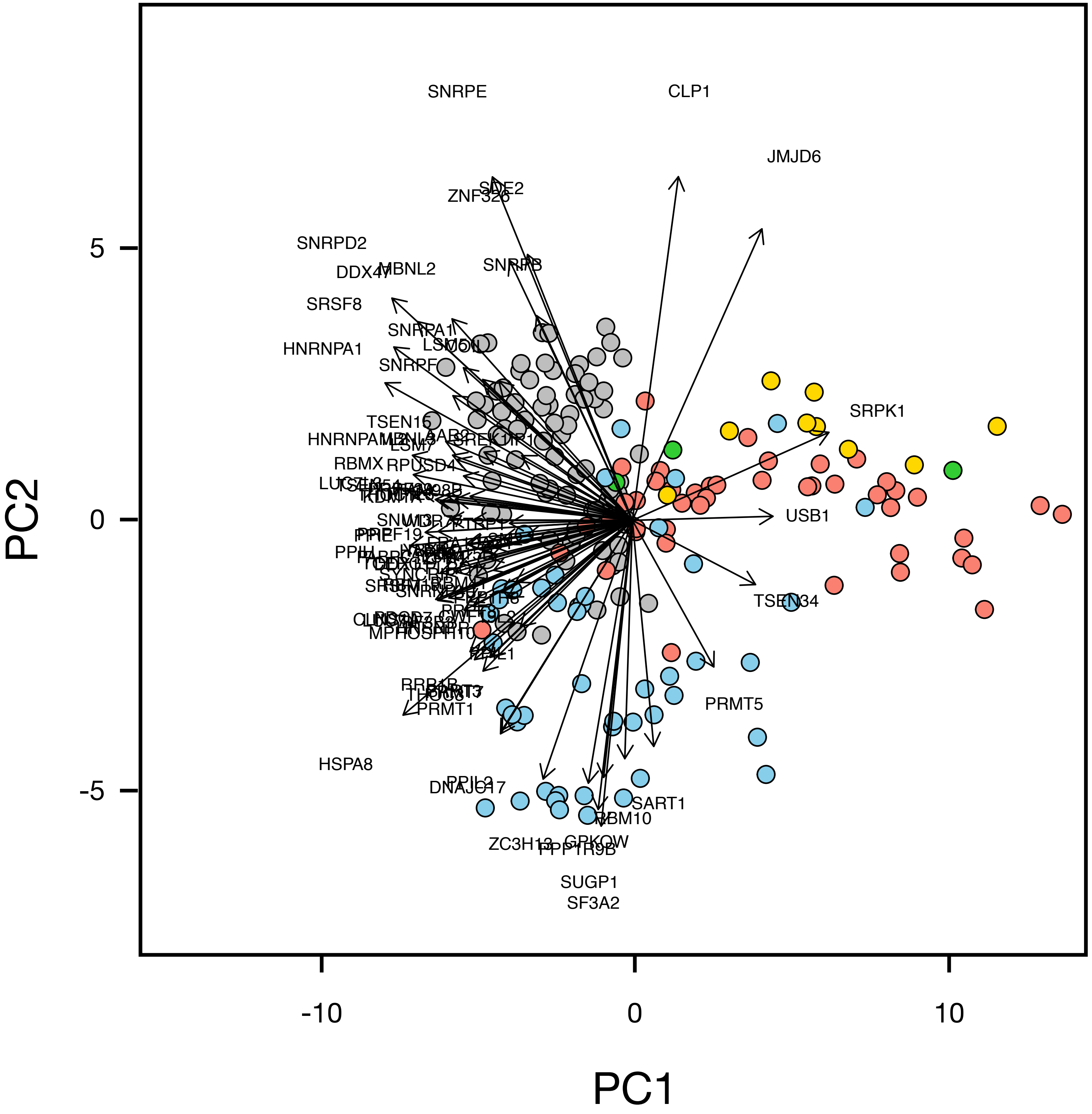


Supplementary figure 3. Original figure of biplot analysis of 304 alternative splicing genes across 190 samples.
